# Exploring the association of collaterals and vessel density using optical coherence tomography angiography in retinal vein occlusions

**DOI:** 10.1101/604983

**Authors:** Hee Eun Lee, Yiyang Wang, Alaa E. Fayed, Amani A. Fawzi

## Abstract

**Purpose:** Using optical coherence tomography angiography (OCTA) to characterize the types of collaterals in eyes with retinal vein occlusion (RVO) and further investigate their correlations with vessel densities of the superficial (SCP) and the deep capillary plexus (DCP).

**Methods:** This cross-sectional study included 25 eyes of 23 patients with RVO. 3 × 3 mm^2^ OCTA macular scans were used to quantify the parafoveal vessel density (VD) of the SCP and DCP, and to classify the collaterals into one of four types (true superficial, true deep, superficial diving, and foveal collateral). Poisson regression model was used to identify significant associations between parafoveal VD and collaterals. We further compared parafoveal VD between subgroups classified by the presence of specific collateral types based on the results of a clustering algorithm.

**Results:** 16 of 25 eyes (64%) developed collaterals. Of the 43 collateral vessels analyzed, 12/19 (63%) true superficial collaterals developed in eyes with central RVO, while all 10 superficial diving collaterals (100%) developed in eyes with branch RVO. Located exclusively in the SCP, true superficial collaterals were all arteriovenous (A-V), while diving collaterals were all veno-venular (V-V). We found a significant negative correlation between SCP VD and the total number of collaterals (R2 = 0.648, P < 0.001) for the entire study cohort. Furthermore, BRVO eyes that developed superficial diving collaterals and CRVO eyes that developed true superficial collaterals demonstrated statistically significant decrease in SCP VD (P-value = 0.014) and DCP VD (P-value = 0.030), respectively, as compared to their counterparts.

**Conclusion:** Our data shows that decreased capillary perfusion in RVO is associated with the development of collaterals, while the RVO type largely dictates the type of collateral that ultimately develops.

## Introduction

Retinal capillary non-perfusion is an important clinical complication of retinal vein occlusion (RVO), which may ultimately compromise the visual prognosis [1, 2]. Driven by hemodynamic factors and increased hydrostatic pressure, collateral vessels may develop secondary to RVO, whereby blood flow surrounding the occluded segment is channeled into areas of lower capillary pressure, opening up collateral vessels [3]. These retinal collateral vessels form within the optic disc or in the retina, appearing as tortuous vessels often crossing the horizontal raphe. Henkind and Wise [4] described collaterals as vessels that originate from existing capillary beds, linking vessels of identical anatomical classification, vein-to-vein in the case of retinal vein occlusion. In their analysis of venous collateral pathways in branch RVO using optical coherence tomography angiography (OCTA), Freund et al [5] found these collaterals to be located exclusively within the deep vascular complex. Distinct from these venous-venous collaterals are those that connect artery to vein, presumably to bypass areas of total capillary bed obstruction. Henkind and Wise differentiated these arterio-venous (A-V) collaterals from vascular shunts that occupy almost the full retinal thickness, forming a direct communication between artery and vein [4].

Our understanding of collateral development is mainly derived from studies in experimental vein occlusion in animal models [6-9]. In human studies, our knowledge of collateral formation is based on RVO natural history studies using fluorescein angiography (FA) [3, 4]. Since FA does not reveal the deep capillaries, or the intraretinal location of collateral vessels, there is a need to better define the patterns of retinal collateral circulation using high-resolution, depth-resolved imaging. Advances in OCTA technology have significantly enhanced our understanding of the connectivity of the human retinal capillary plexuses [10-13]. Driven by the greater axial resolution of this technology, studies have shown that individual capillary plexuses may be disproportionately affected in retinal vascular disease [14, 15].

The impact of collateral vessels on the prognosis and management in RVO is still a matter of ongoing debate. At one end of the spectrum, collaterals have been hypothesized to provide an alternative source of drainage for the impaired retinal circulation [2] and have been reported to predict good visual prognosis [16, 17]. Using OCTA, Heiferman et al demonstrated that the path of perfused collaterals in eyes with RVO showed relatively preserved focal inner retinal structure despite the large zones of surrounding capillary non-perfusion [18]. Their findings suggest that perfused collaterals on OCTA may mitigate ischemic retinal atrophy by providing focal inner retinal perfusion in these ischemic zones.

Visual acuity improvement after the resolution of hemorrhage and edema is often limited by the extent of capillary non-perfusion at the macula in eyes with RVO [19]. Understanding whether the perfusion status of the retinal vascular plexuses is related to collateral vessel formation could provide important insights regarding the prognostic value of collaterals in RVO. To address this gap in knowledge, we designed the current study to characterize macular collaterals in RVO and to investigate the correlation between their development and capillary vessel density in the macular superficial and deep plexuses.

## Methods

This was a cross-sectional analysis of RVO patients who underwent OCTA in the Department of Ophthalmology at Northwestern University of Chicago, Illinois between December 2, 2015 and July 23, 2018. This study was approved by the Institutional Review Board of Northwestern University, followed the tenets of the Declaration of Helsinki and was performed in accordance with the Health Insurance Portability and Accountability Act regulations. Written informed consent was obtained from all participants.

### Study Sample

Inclusion criteria for this study required a diagnosis of retinal vein occlusion based on the clinical assessment by one experienced retinal specialist (A.A.F.). The three main types of RVO included were central (CRVO), branch (BRVO), and hemi-central retinal vein occlusion (HRVO). The study included treatment-naïve eyes and those treated previously with anti-vascular endothelial growth factor (VEGF) and laser. Patients with RVO involving the macula and temporal sector were included. However, patients with RVO in which the occluded site was located in the nasal sector were excluded. FA images were examined to define the location of vein occlusion. In four eyes, FA was not available at the time of examination and spectral-domain OCT (Heidelberg Spectralis HRA + OCT; Heidelberg Engineering, Heidelberg, Germany) was used instead to define areas of occlusion. Additionally, eyes with coexisting ocular diseases such as retinal arterial occlusion and diabetic retinopathy were excluded. We also excluded eyes for which inadequate quality OCTA images were obtained due to eye movement or cataract, and a signal strength index (SSI) score below 50. At initial visits, patients underwent fundus biomicroscopy and a comprehensive bilateral ophthalmic examination including FA and SD-OCT.

### OCT Angiographic Imaging

RTVue-XR Avanti system (Optovue Inc., Fremont, California, USA) was used to acquire a 3 × 3 mm^2^ scanning area, centered on the fovea. RTVue-XR Avanti system captures at each location two consecutive B-scans (M-B frames), with each B-scan containing 304 A-scans. The A-scan rate is 70,000 scans per second and uses a light source centered on 840 nm and a bandwidth of 45 mm. OCTA incorporates the split-spectrum amplitude-decorrelation angiography (SSADA) software (version 2017.1.0.151) [20] to extract angiographic flow information by using an algorithm that quantifies the decorrelation of the OCT reflectance between the two consecutive B-scans.

### Image analysis, OCT Angiographic Evaluation of Parafoveal Vessel Density and Classification of Collateral Vessels

OCTA images taken on each patient’s follow-up visit were used to examine the incidence of collateral vessels in the macular area. We chose to analyze OCTA scans from the last visit for patients who underwent multiple imaging sessions coinciding with follow-up visits, provided that a sufficient time (> 6 months) has elapsed since RVO onset; most collaterals have been reported to develop > 6 months after disease onset [2], though some develop within 6 months [3]. If a patient had multiple OCTA scans for the follow-up visit, OCTA scan with the better image quality was chosen for analysis. In the 3 × 3 mm^2^ OCTA image, the parafoveal vessel density (VD) of the SCP and DCP was quantified using the AngioVue Analytics software (version2017.1.0.151) of the OCTA device. The software calculates the area occupied by blood vessels within a ring-shaped region of interest centered on the fovea with an inner and outer ring diameter of 1 mm and 3mm, respectively. The parafoveal VD is automatically reported as a percentage of the total area.

The built-in software of the OCTA device automatically defines the segmentation boundaries used to visualize SCP, DCP, and the full retinal thickness. The SCP slab was segmented from the internal limiting membrane (ILM) to 9 μm above the inner plexiform layer (IPL). The DCP slab was segmented from 9 μm above the IPL to 9 μm below the outer plexiform layer (OPL). The built-in software sets the inner boundary of full retinal slab at the ILM and the outer boundary at 9 μm below the OPL. In cases of automated software segmentation error due to macular edema or macular thinning, manual adjustment of the segmentation boundaries was performed. Eyes were classified as having macular edema secondary to RVO if they had a central macular thickness (CMT) of equal to or greater than 300 μm [21]. CMT was also recorded using the same built-in software.

All collateral vessels were identified within the 3 × 3 mm^2^ en face angiograms of each eye. Images were further analyzed to assess the location of each collateral vessel using the corresponding cross-sectional scans with angiographic flow overlay, and 3D Projection artifact removal (PAR) technology to eliminate projection artifact. We excluded non-perfused collateral vessels that were visualized on en *face* OCT without corresponding flow detected on OCTA. The collaterals selected for analysis were classified into one of four types of collaterals (true superficial, true deep, superficial diving, and foveal collateral) based on whether they coursed through the SCP or DCP, and the location of parent vessels connected to the collateral itself, as depicted by Fig 1 and 2. Collaterals connecting parent blood vessels in the SCP and those connecting parent blood vessels in the DCP were defined as true superficial collaterals and true deep collaterals, respectively. Collaterals connecting two superficial vessels via the DCP were classified as superficial diving collaterals. Foveal collaterals are defined as collaterals that cross the horizontal raphe across the fovea and connect two superficial blood vessels. Since the segmentation of the SCP terminates closer to the fovea, foveal collaterals are distinguished from the superficial diving collaterals. The collaterals were further classified into arterio-venous (A-V) or veno-venular (V-V) using color fundus photographs and FA images. We differentiated A-V collaterals from shunt vessels by ensuring that the selected vessels were not occupying the entire retinal thickness or protruding from the retinal surface [4]. Since we evaluated the identity of the parent vasculature (arterioles vs. venules) using FA, we could only classify collaterals connecting parent vessels in the SCP into A-V or V-V collaterals.

**Fig 1.**
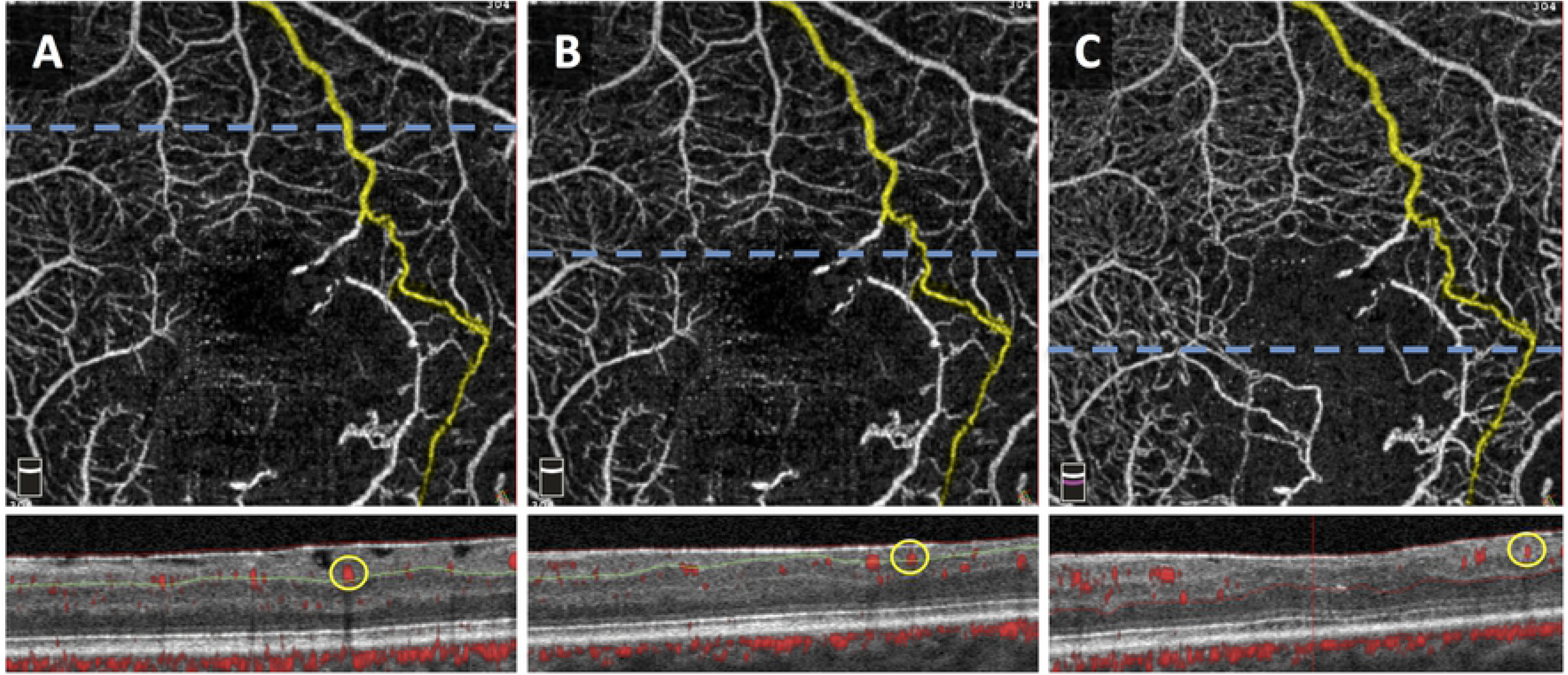
3 × 3-mm^2^ *en face* Optical Coherence Tomography Angiography (OCTA) images of the same eye with a true superficial collateral crossing the horizontal raphe. Segmentation of the SCP (A and B) and full retinal slab (C) illustrate that the collateral is confined to the SCP. Horizontal (blue) dashed lines correspond to OCT B-scans with red flow overlay shown below each *en face* image. The circle (yellow) in the B-scans corresponds to the flow signal of the collateral vessel, confirming its location in the SCP. Green and red lines in B-scan images demonstrate segmentation boundaries for the SCP and full retina, respectively.

**Fig 2.**
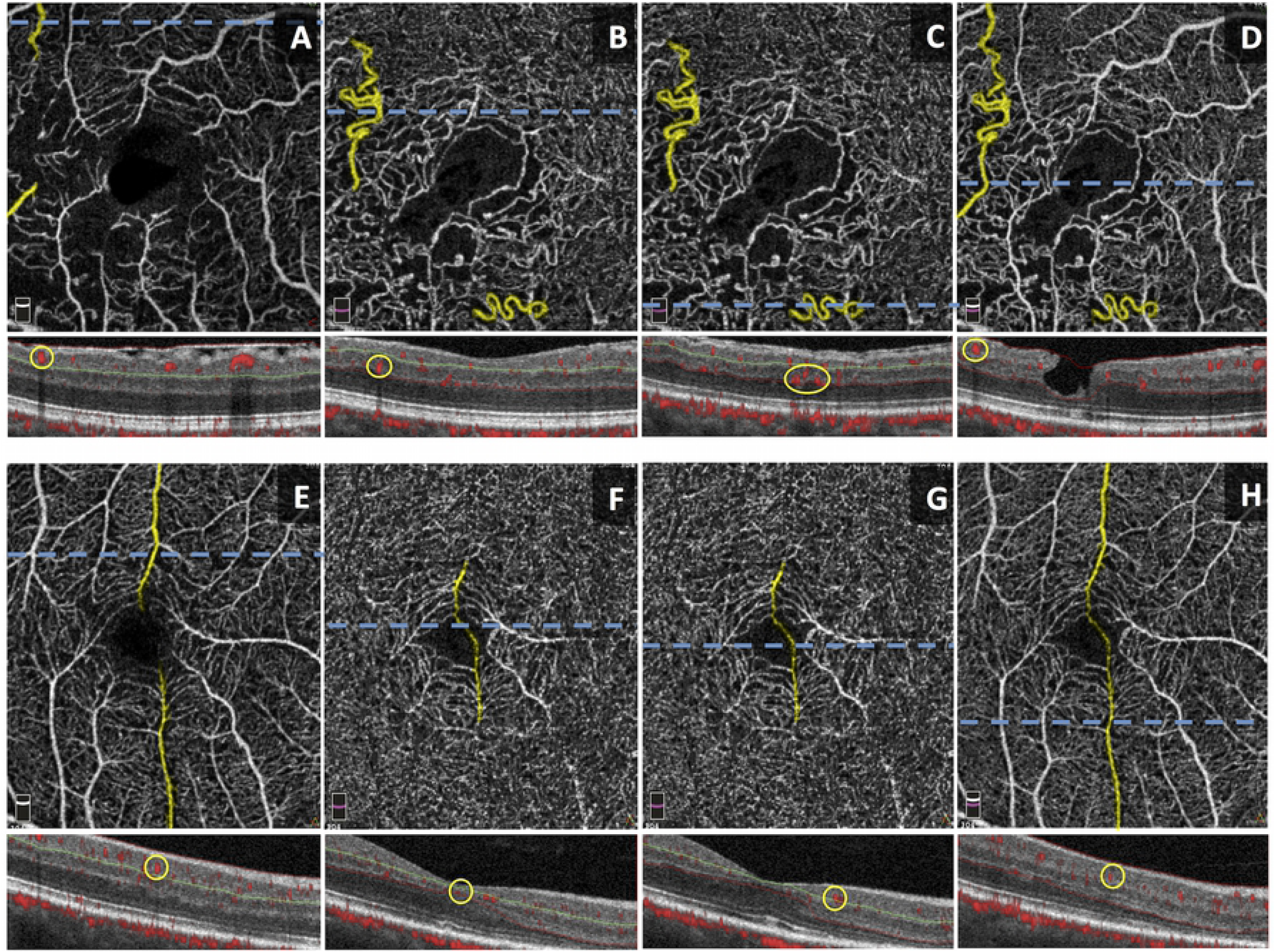
Optical Coherence Tomography Angiography (OCTA) Imaging of Collaterals. Top row (A-D) shows OCTA *en face* images of an eye with both superficial diving and true deep collaterals, and corresponding OCT B-scans with red flow overlay directly below. Segmentation of the SCP (A) and DCP (B and C) illustrate that the superficial diving collateral connects its parent vessels in the SCP by coursing through the DCP, whereas the true deep collateral is confined to the DCP. The full retinal thickness slab shows the combined superficial diving collateral and its parent vessels in one segmentation. Bottom row (E-H) shows OCTA *en face* images of an eye with a foveal collateral, and corresponding OCT B-scans with red flow overlay directly below. Segmentation of the SCP (E) and DCP (F and G) illustrate two parent vessels connected at the fovea by the foveal collateral (H). Horizontal (blue) dashed lines correspond to OCT B-scans with red flow overlay shown below each *en face* image. The circle (yellow) in the B-scans correspond to the flow signal of the collateral vessel. Green lines in B-scan images demonstrate segmentation boundary between the SCP and DCP.

### Statistical Analysis

To better understand the patterns in our data, we implemented the Partitioning Around Medoids (PAM) clustering algorithm [22] by dividing the data (Table 1) into distinct groups using the selected features: “RVO Type”, “LogMAR BCVA”, “SCP VD” and “DCP VD”. Using this algorithm, it is expected that similar patterns will be observed within the clusters and different patterns between the clusters. Since “RVO Type” in Table 1 is a categorical feature while the other features implemented into the algorithm are numerical, we adopted the Gower Distance [23] as a similarity measurement between the two types of features. For the categorical feature, we first converted the 3 RVO types (BRVO, CRVO, and HRVO) into 3 binary columns then calculated the Sørensen–Dice index distance [24, 25]. On the other hand, we normalized the numerical features using the Min-Max normalization method before calculating the Manhattan Distance [26]. The Gower Distance between these two data records is the average distance between all features. PAM clustering algorithm also requires an initial number of clusters, k. We varied k from 2 to 10 and calculated the Silhouette Width value [27] for each k value; we chose the k value with the maximum Silhouette Width value. The steps in the procedure of PAM Clustering Algorithm are as follows:

**Table 1.**
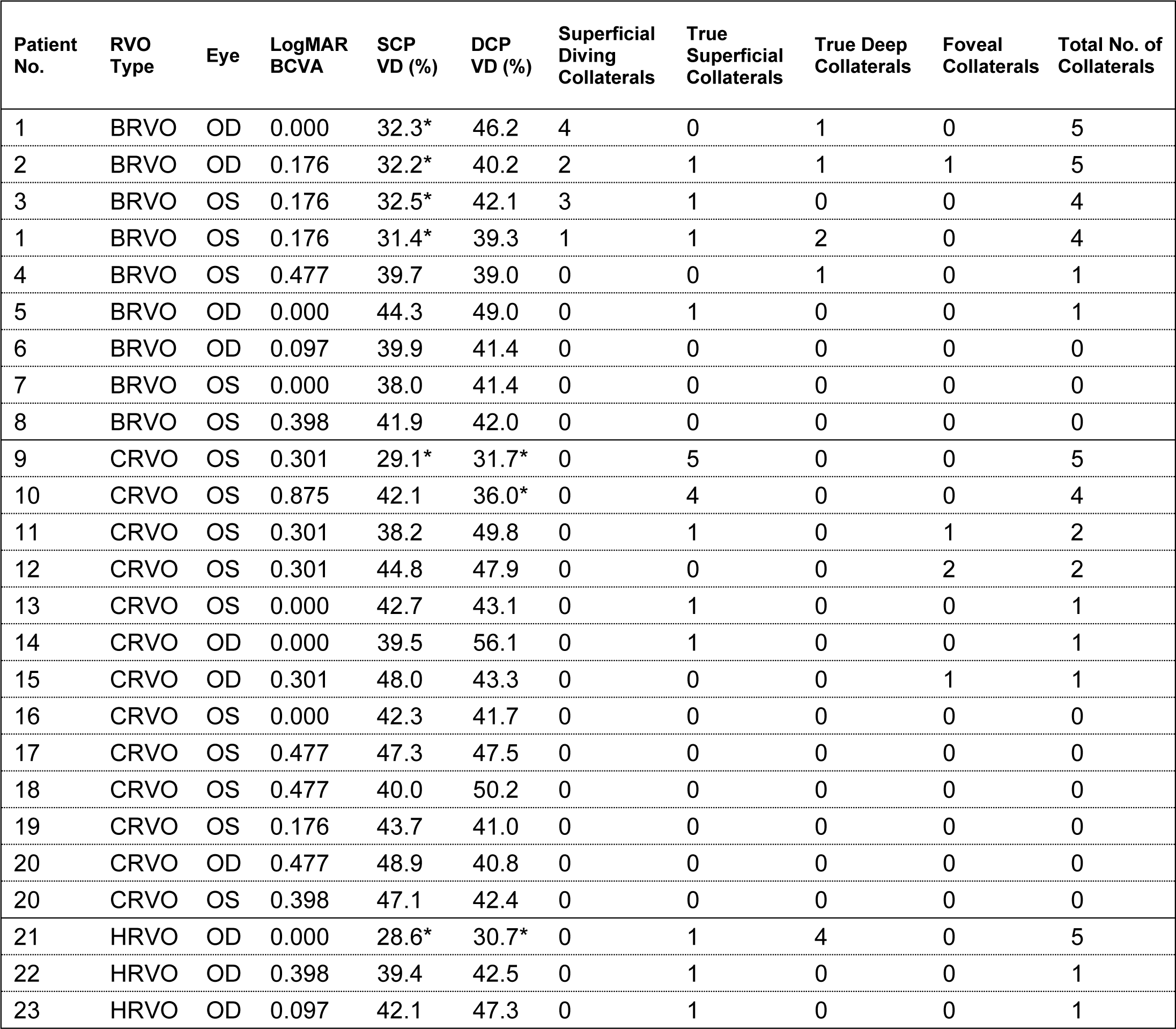
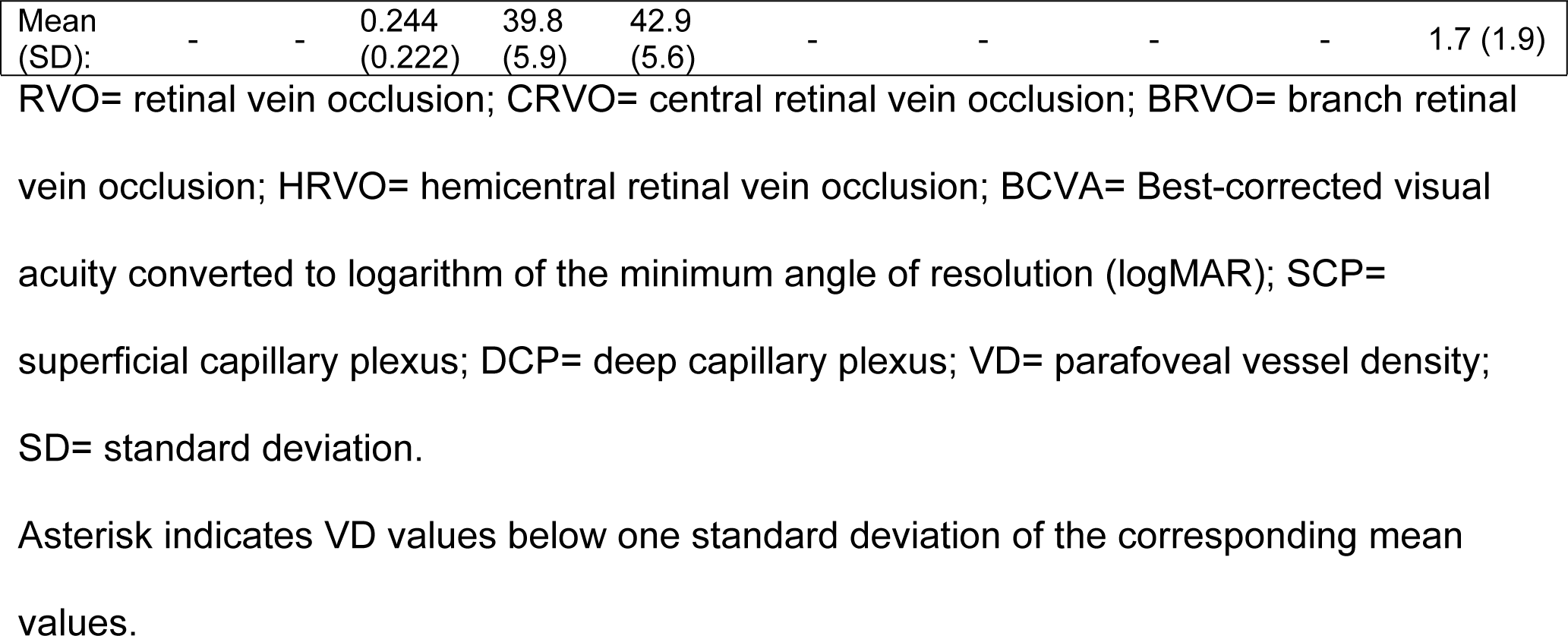
Number of Collateral Vessels for Each Study Eye in 3 × 3-mm^2^ Optical Coherence Tomography Angiography Scans

1. Randomly choose k data records as the medoids
2. For every data record, find and group with its closest medoid
3. For each cluster, find the data record with the lowest average distance to the rest of the data records within the cluster. If the data record is different from the previous medoid, replace the previous medoid with the data record as the new medoid.

Any changes to the medoids in this procedure will cause the algorithm to repeat Step 2. If all the medoids remain unchanged, the algorithm will end the procedure.

We used a Poisson regression model to investigate the associations between the number of collaterals and OCTA parameters (SCP and DCP VD) for the entire study cohort. Exploratory data analysis indicated non-normality of the collateral count data; hence, Poisson regression model was the optimal fit for our small sample size. For subgroup analyses, we used the Mann-Whitney U, non-parametric test to compare differences in SCP and DCP VD observed in RVO types classified by the presence of specific collateral types (superficial diving collaterals in BRVO and true superficial collaterals in CRVO), as revealed by the clustering results. All statistical analyses were conducted using IBM SPSS statistics version 25 (IBM SPSS Statistics; IBM Corporation, Chicago, IL, USA). P < 0.05 was considered to be the threshold for statistical significance.

## Results

Between December 2, 2015 and July 23, 2018, 40 eyes of 38 patients with RVO underwent imaging with OCTA at Northwestern University. Of these eyes, 14 were excluded from the study due to insufficient image quality. One eye had a superonasal RVO outside the macular scan and was further excluded from the analysis. Of the remaining 25 eyes of 23 subjects, nine eyes (36%) had BRVO, 13 eyes (52%) had CRVO and three eyes (12%) had HRVO. Overall, 16 of 25 eyes (64%) developed collaterals; 6/9 eyes with BRVO, 7/13 eyes with CRVO and 3/3 eyes with HRVO developed collaterals. Patient characteristics are summarized in Table 2.

**Table 2.**
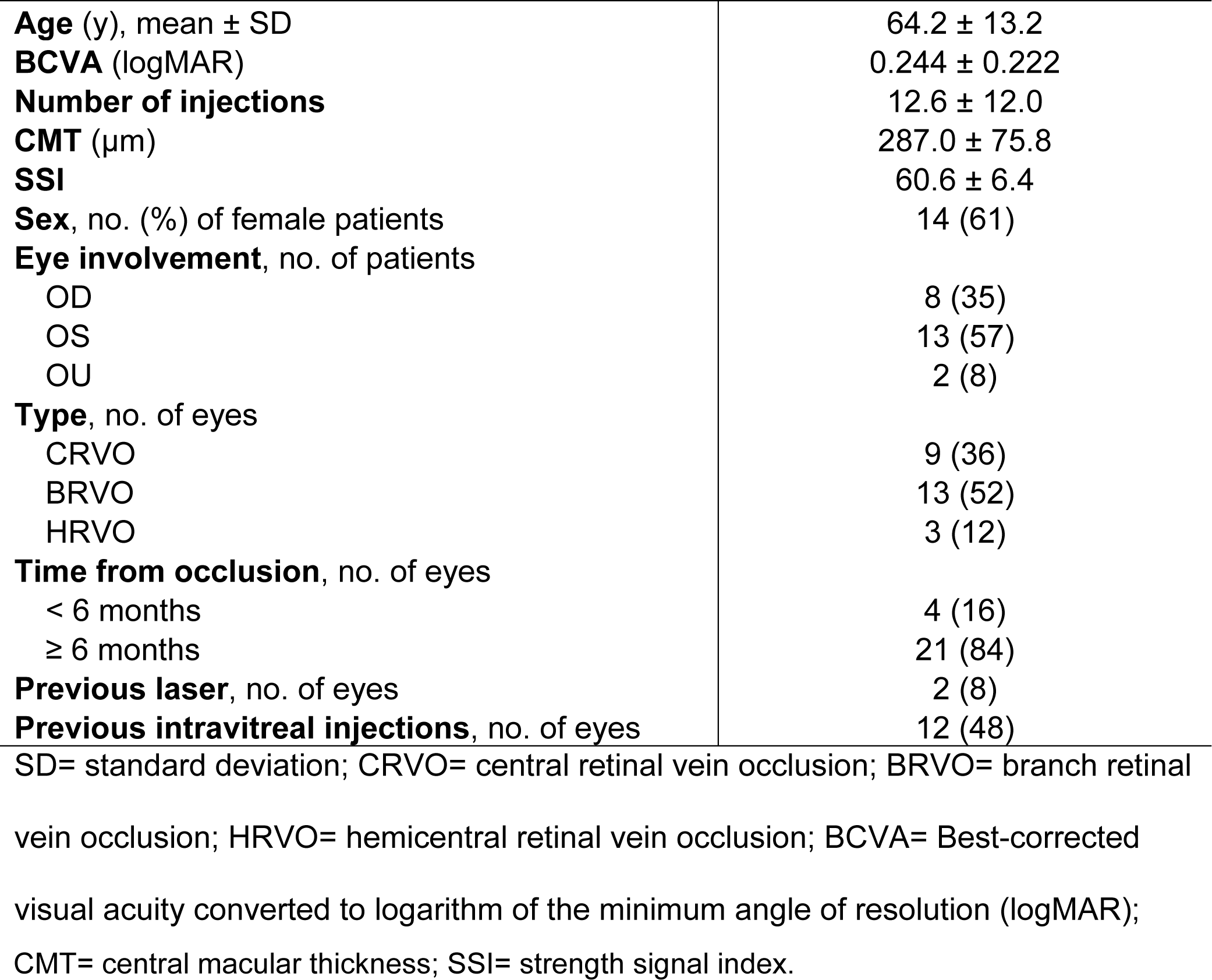
Characteristics of 23 Patients (n = 25 eyes) with Retinal Vein Occlusion at the Time of Imaging with Optical Coherence Tomography Angiography

### Image analysis, OCT Angiographic Evaluation of Parafoveal Vessel Density and Classification of Collateral Vessels

The mean parafoveal VD (%) in the SCP and DCP within a 3 × 3-mm^2^ area were 39.8 ± 5.6 and 42.9 ± 5.6, respectively. SCP VD or DCP VD values below one standard deviation of their corresponding mean values were considered “low”. 43 collateral vessels were identified and analyzed (mean, 1.7; range, 0-5 collaterals per eye). We found that the number of collaterals was highest (≥ 4 collaterals per eye) in eyes with either low SCP or DCP VD (Table 3).

**Table 3.** Optical Coherence Tomography Angiography Findings in 3 × 3-mm^2^ Scans (n = 25 eyes) in Eyes with Retinal Vein Occlusion

|  | **CRVO (n = 9)** | **BRVO (n = 13)** | **HRVO (n = 3)** |
| --- | --- | --- | --- |
| **Collaterals, no. (%)** |  |  |  |
| True superficial | 12 (63) | 4 (21) | 3 (16) |
| True deep | 0 (0) | 5 (56) | 4 (44) |
| Superficial diving | 0 (0) | 10 (100) | 0 (0) |
| Foveal | 4 (80) | 1 (20) | 0 (0) |
| **Collaterals, no. of eyes** |  |  |  |
| A-V | 5 | 1 | 2 |
| V-V | 0 | 3 | 1 |
| **Collateral Group (n = 16)** |  |  |  |
| SCP VD (%), mean ± SD | 40.6 ± 6.0 | 35.4 ± 5.3 | 36.7 ± 7.1 |
| DCP VD (%), mean ± SD | 44.0 ± 8.3 | 42.6 ± 4.1 | 40.2 ± 8.5 |
| **Non-Collateral Group (n = 9)** |  |  |  |
| SCP VD (%), mean ± SD | 44.9 ± 3.4 | 40.0 ± 2.0 | - |
| DCP VD (%), mean ± SD | 43.9 ± 3.9 | 41.6 ± 0.3 | - |

CRVO= central retinal vein occlusion; BRVO= branch retinal vein occlusion; HRVO=
hemicaltral retinal vein occlusion; A-V= arterio-venous; V-V= veno-venular (venous);
SCP= superficial capillary plexus; DCP= deep capillary plexus; VD= parafoveal vessel
density; SD= standard deviation.

We examined the overall mean VD of the RVO types, classified by presence of collaterals, as shown in Table 3. The mean DCP VD was similar for CRVO and BRVO eyes, regardless of the presence of collaterals. Interestingly, the mean SCP VD was lower for both CRVO and BRVO eyes with collaterals. Poisson regression model (S1 Table) showed that SCP VD was most significantly associated with the total number of collaterals (P < 0.001). The exponentiated value of coefficient estimate was 0.86, indicating that the incidence rate of collaterals will increase by 14% for every unit decrease in SCP VD. According to the results of the subgroup analyses using the Mann-Whitney U test, 4 eyes with BRVO (Patient no. 1-3, Table 1) that developed superficial diving collaterals had significantly lower SCP VD values than the remaining BRVO eyes (P = 0.014). In addition, 2 eyes with CRVO (Patient no. 9-10, Table 1) that developed the highest number of true superficial collaterals (≥ 4 collaterals per eye) had significantly lower DCP VD values than other eyes with CRVO (P = 0.030).

In the classification of collateral vessels into A-V or V-V collaterals, 4 of the 16 eyes that formed collateral vessels did not have adequate fundus photographs or FA images for analysis. All eyes with BRVO that developed collaterals had V-V collaterals, while all of the eyes with CRVO had A-V collaterals and none had V-V collaterals (Table 3).

### Cluster analysis

Using the PAM clustering algorithm, the number of clusters, k, was determined to be 3, based on the calculated maximum Silhouette Width value. The algorithm divided the eyes into 3 separate clusters based on the implemented features, including VD of the SCP and DCP. Each cluster was homogeneous for RVO type (CRVO, BRVO, and HRVO). The CRVO cluster had the highest mean SCP VD and DCP VD, and the most foveal and A-V collaterals. The HRVO cluster had the highest mean value of true deep collaterals. Both CRVO and HRVO clusters had a high mean value of true superficial collaterals. The BRVO cluster had the highest mean value of diving collaterals and the most V-V collaterals.

## Discussion

Our study shows that the number of collaterals is significantly associated with the extent of capillary non-perfusion in RVO, suggesting that collateral formation may be a potential marker of retinal ischemia. In a natural history study of non-ischemic CRVO eyes that developed collaterals (n=174), 52% had 20/70 or worse visual acuity compared to 35% in eyes that did not develop collaterals (n=109) [28]. The authors suggested that the poor visual acuity was due to the fewer available tributaries in patients that developed collaterals. This hypothesis may be supported with our finding that with decreasing capillary density, the number of collaterals increases (Table 1). The correlation between extent of collaterals and visual prognosis remains to be validated in future studies.

While retinal ischemia is significantly associated with collateral formation in RVO, our data suggests that it is the RVO type that ultimately dictates the specific type of collateral. We used an unsupervised clustering algorithm in order to further investigate whether SCP and DCP densities alone could differentiate collateral types. We found that collaterals fell into 3 distinct groups, largely distinguished by the RVO type (CRVO, BRVO, and HRVO), rather than vessel density. Clustering revealed a similarity in collateral types within each of these groups, with true superficial collaterals (Fig 1) occurring predominantly in the CRVO and HRVO groups, while the superficial diving collaterals (Fig 2) were found in the BRVO group. Furthermore, multivariate analysis of the entire study cohort showed that non-perfusion at the SCP capillary plexus in RVO was most significantly associated with the total number of collaterals (P < 0.001). The apparent lack of similar association of DCP VD could be due to the heterogeneity of our study patients including the type of occlusion present (BRVO, CRVO, and HRVO) [29] that may have introduced variations to the regression model (S1 Table). It is possible that with a greater number of cases, a stronger correlation between DCP VD and the overall number of collaterals in RVO could be revealed.

When we divided the eyes into RVO subgroups based on the aforementioned clustering results, BRVO eyes that developed superficial diving collaterals had significantly lower SCP VD (P = 0.014, Mann-Whitney U test) than those that did not. It is possible that diving collaterals form as a secondary complication in response to primary SCP capillary closure. In this model, diving collaterals (Fig 2A-2D) would serve as a channel for SCP venous outflow, bypassing the occluded SCP capillary bed, with the DCP providing a more convenient outflow through its highly connected vortex capillary system that naturally crosses the horizontal raphe [10]. Supplementary file (S2 Video) shows a video fly-through revealing a superficial collateral diving into the DCP. A prospective longitudinal study is needed to confirm that collaterals indeed form as a result of SCP capillary loss.

Wakabayashi and associates have shown that preserved perfusion in the DCP is associated with more favorable visual acuity in BRVO [19]. Ischemia in the deep vascular network is commonly associated with macular edema [30, 31], and collaterals coursing through the DCP can possibly improve visual outcome by providing a drainage pathway [5]. It is possible that the superficial diving collaterals in BRVO eyes observed in our study may be associated with better perfusion in the DCP leading to better visual acuity. Kang et al reported that the visual acuity is positively correlated with the deep parafoveal vessel density in 33 eyes with RVO [32]. These results were recently corroborated by Seknazi et al in a study of 65 eyes with RVO [33]. Taken together, these studies suggest that visual acuity reduction may be related to decreased DCP vessel density. The ability of diving collaterals to improve DCP perfusion and consequently, achieve a positive visual prognosis needs to be further explored. We were unable to investigate the relationship between collateral formation and visual acuity due the small sample size and the presence of other confounding factors that could contribute to visual outcomes in our current study, including cataract surgery and anti-VEGF treatment status.

According to Henkind and Wise [4], macular venous-venous (V-V) collaterals are more commonly seen in patients with BRVO, while in CRVO these collaterals develop at or within the optic disc. This is in line with our finding that macular collaterals encountered in eyes with CRVO were all A-V collaterals, while in eyes with BRVO the majority were V-V (Table 3). Optic nerve V-V collaterals in eyes with CRVO were most likely located outside the boundaries of the 3 × 3 mm^2^ OCTA scan area. None of the V-V collaterals in BRVO were located in the SCP, consistent with the recent findings of Freund et al [5]. In addition to their findings, our study explored A-V collaterals, which were confined to the SCP in CRVO (n = 12/12 true superficial collaterals). The A-V collaterals are postulated to form in situations where capillary beds are completely obstructed in order to allow blood to cross from the arterial to the venous side, as shown by fluorescein angiography studies [4, 34]. However, without the dampening effect of capillaries, these collaterals have lower resistance to blood flow and can therefore paradoxically induce ischemia in the surrounding retina [35]. In our study, the mean SCP VD was lower in CRVO eyes that developed A-V collaterals (Table 3). Therefore, it is likely that decreased perfusion observed in these CRVO eyes may be related to the preferential flow of blood through these low resistance, high flow channels in A-V collaterals [36].

A number of studies have reported that collateral development in RVO is not influenced by therapeutic interventions [37-39]. The development of collaterals in BRVO was not significantly different comparing eyes that received either anti-VEGF or grid laser therapy [37, 40]. More recently, Falavarjani et al showed that anti-VEGF injections did not affect SCP or DCP perfusion densities [41]. It is therefore not certain that treatments such as anti-VEGF injections or laser therapy in our study could have contributed to capillary loss or collateral vessel formation.

Strengths of our study include the stringent criteria for quality of our datasets, including high-signal strength index, 3 × 3-mm^2^ OCTA scans which made it possible to obtain the most accurate of VD in the macula at the different plexuses [42, 43]. Due to the limitation of the scan size, however, our analysis did not include the peripheral collaterals. In addition, we carefully assessed the OCTA cross-sectional scans to trace the flow signals of the collateral vessels in the *en face* images (Fig 1 and 2) and performed manual adjustment to correct OCTA segmentation in eyes with macular edema or retinal thinning. Being a cross-sectional study based on a small number of patients, this study has apparent limitations. The study included a limited sample size of HRVO patients (n = 3), which may have contributed to the lack of significant findings in this type of RVO. Although the current study only included perfused collaterals with flow signal that represented more physiologic levels of retinal blood flow, discordance between flow on OCTA and vasculature on *en face* OCT have been previously reported [18], emphasizing the fact that OCTA technology may simply lack the sensitivity to detect sub-threshold blood flow in some collaterals.

In summary, our cross-sectional study of 43 collateral vessels in 25 eyes with RVO showed that vessel density in superficial and deep capillary networks is significantly associated with the development of collaterals. Our clustering algorithm suggests that specific collateral types develop in different RVO types [5, 29], with the addition of an improved understanding of the particular role played by perfusion defects in the SCP and DCP. Further studies in larger cohorts are required to test the hypothesis that baseline perfusion status of the retinal capillary layers to predict collateral formation in RVO, in addition to investigating the correlation between collaterals, perfusion and visual function.

